# Restoration of functional PAX6 in aniridia patient iPSC-derived ocular tissue models using repurposed nonsense suppression drugs

**DOI:** 10.1101/2022.10.12.511600

**Authors:** Dulce Lima Cunha, Jonathan Eintracht, Philippa Harding, Jo Huiqing Zhou, Mariya Moosajee

## Abstract

Aniridia is a rare, pan-ocular disease causing severe sight loss, with only symptomatic intervention offered to patients. Approximately 40% of aniridia patients present with heterozygous nonsense variants in *PAX6*, resulting in haploinsufficiency. Translational readthrough inducing compounds (TRIDs) have the ability to weaken the recognition of in-frame premature stop codons (PTCs), permitting full-length protein to be translated. We have established induced pluripotent stem cell (iPSC)-derived 3D optic cups and 2D limbal epithelial stem cell (LESC) models from an aniridia patient with a prevalent *PAX6* nonsense mutation. Both *in vitro* models show reduced PAX6 protein levels, mimicking the disease. Repurposed TRIDs amlexanox and 2,6-diaminopurine (DAP), and positive control compounds ataluren and G418 were tested for their efficiency. Amlexanox was identified as the most promising TRID, increasing full-length PAX6 levels in both models, and rescuing the disease phenotype through normalization of VSX2 and cell proliferation in the optic cups and reduction of ABCG2 protein and *SOX10* expression in LESC. This study highlights the significance of patient iPSC-derived cells as a new model system for aniridia and proposes amlexanox as a new putative treatment for nonsense-mediated aniridia.

## Introduction

Aniridia (OMIM 106210) is a rare, dominant, pan-ocular disease, with a prevalence of 1 in 40,000-100,000 ^1^. Typical symptoms of this disease include congenital iris and foveal hypoplasia with nystagmus, and progressive development of glaucoma, cataracts and keratopathy, leading to significant visual impairment ^2,3^. Up to 90% of aniridia patients develop limbal stem cell deficiency (LSCD), where adult epithelial stem cells originating in the limbus and maintain corneal transparency, are lost or defective, causing impaired epithelium renewal and conjunctival invasion ^4^. LSCD invariably results in complete corneal opacity, usually termed aniridia-related keratopathy (ARK), and is the most relevant feature contributing to visual loss in aniridia post-natally ^4,5^.

Heterozygous mutations affecting the *PAX6* gene or its regulatory regions are the cause of aniridia ^6,7^, with mutations introducing a premature stop codon (PTC) being the most common (http://lsdb.hgu.mrc.ac.uk/home.php?select_db=PAX6). Of these, nonsense mutations are the most prevalent, accounting for 39% of the total mutations reported in aniridia patients ^8^. *PAX6* nonsense mutations are predicted to result in loss of function, where mutated mRNA is likely degraded by nonsense-mediated decay (NMD), resulting in *PAX6* haploinsufficiency.

Nonsense suppression or translational readthrough inducing compounds (TRIDs) weaken the recognition of a PTC and promote the replacement of a near cognate amino acid, thus allowing translation to continue and producing a full-length protein ^9,10^. Promising preclinical data using ataluren (also called Translarna or PTC124), a TRID approved for the treatment of Duchenne muscular atrophy, showed rescue of Pax6 levels in the aniridia *Sey*^+/−^ mouse model, with topical administration inhibiting disease progression and improving corneal, lens, and retinal defects ^11,12^. A phase I/II clinical trial (NCT02647359) for aniridia was completed, but failed to meet the primary endpoint, despite showing a positive trend towards functional improvement (https://www.prnewswire.com/news-releases/ptc-therapeutics-reports-fourth-quarter-and-full-year-2019-financial-results-and-provides-a-corporate-update-301014669.html). The use of TRIDs is a particularly suitable therapeutic approach for aniridia due to the high prevalence of nonsense variants and the milder phenotype associated with *PAX6* missense mutations ^2,3,8^. However, novel readthrough compounds with improved efficiency are required with a personalized medicine approach, knowing which TRIDs may be more effective for specific PTCs or that combined inhibition of nonsense-mediated decay may boost mRNA substrate and end protein production.

Pax6 is a dose-sensitive transcription factor essential for eye development ^6,7^. It is expressed early in ocular morphogenesis, during the establishment of the eye field and optic vesicle, and has multiple roles in the development and maintenance of retinal progenitor cells, lens, cornea and iris ^13^. In the cornea, correct Pax6 levels are required for normal cell growth during limbal and central corneal epithelial development, but the exact mechanisms on how *PAX6* haploinsufficiency causes LSCD and ARK are still not understood ^14^. It was recently shown that PAX6 controls neural crest migration during corneal development, a process important for the formation of the non-epithelial corneal layers, i.e. stroma and endothelium, as well as for maintenance of the limbal niche ^15–17^.

The generation of human induced pluripotent stem cells (iPSCs) has opened a new avenue in establishing representative *in vitro* models that can recapitulate human development and provide valuable insights on disease mechanisms ^18^. They have been used to accelerate therapeutic development in several retinal and corneal eye disorders ^19–21^. This is the first study to generate iPSCs from an aniridia patient carrying a heterozygous *PAX6* nonsense mutation with a UGA-type PTC and establish patient-specific iPSC-derived optic cups and limbal epithelium stem cell (LESC) models that mimic the haploinsufficiency state. We used these models to assess the potential of TRIDs amlexanox and 2,6-diaminopurine (DAP) to treat aniridia. Amlexanox is a FDA-approved drug used for the treatment of asthma and aphthous mouth ulcers ^22^, that was found to have both readthrough and NMD-inhibition properties ^23–25^; DAP is an antileukemia compound with recently identified strong readthrough capacity for UGA-type PTCs ^26^.

We identified amlexanox as the most promising TRID, increasing full-length PAX6 levels and rescuing phenotype abnormalities in both iPSC-derived retinal and corneal models, while DAP showed distinct tissue dependent responses. Our results provide substantial proof-of-concept for the use of amlexanox as a new therapeutic approach for aniridia.

## Results

### Generation of aniridia induced pluripotent stem cells (iPSCs)

Human dermal fibroblasts taken from a molecularly confirmed aniridia patient (AN) were reprogrammed into iPSCs by electroporation using non-integrating episomal plasmids ^27,28^. Generated iPSC clonal lines were thoroughly and routinely characterized, showing positive pluripotency markers, tri-lineage differentiation ability and chromosomal stability (Supp Figure 1). AN patient carries a pathogenic heterozygous nonsense variant in *PAX6:* c.781C>T/ p.(Arg261*) (NM_000280.4) ^8^. The disease-causing variant was confirmed in AN iPSCs by direct sequencing of *PAX6* exon 10 (Figure 1A).

**Figure 1.**
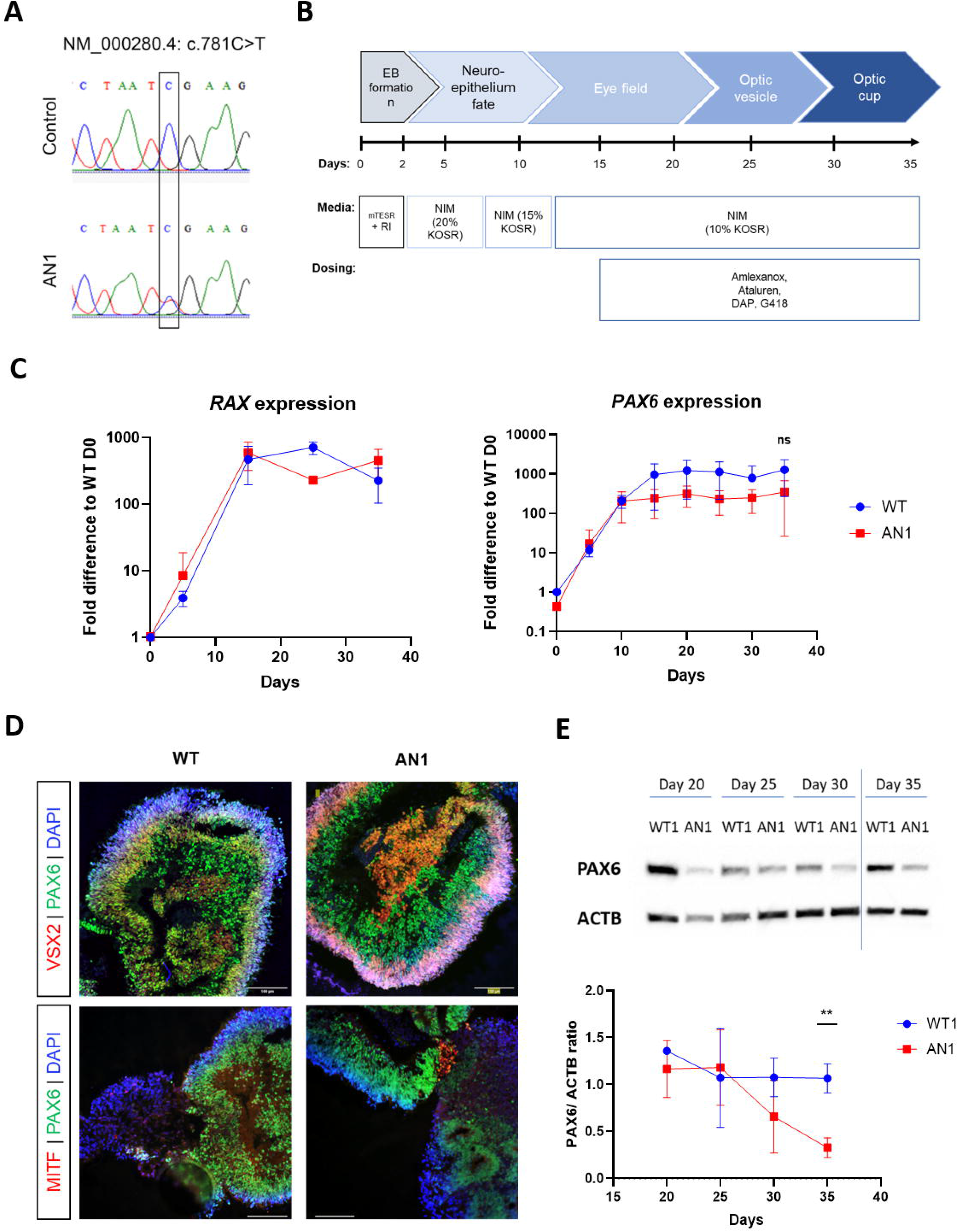
Generation of aniridia iPSC-derived optic cups. **(A)** Direct sequencing of *PAX6* exon 10 showing the heterozygous nonsense c.781C>T change in aniridia patient iPSCs. This variant was not detected in control lines. **(B)**Schematic representation of the differentiation strategy of iPSCs into 3D optic cups (35 days). Dosing experiments with TRIDs were performed from day 15 onwards. **(C)** qRT-PCR transcript analysis of eye field transcription factors *RAX* and *PAX6* during 35 days of differentiation in control (WT, blue) and aniridia (AN, red) iPSCs. Values were normalised to day 0 and to internal housekeeping gene *GAPDH*. Data represent means and SD of n=3 biological replicates. **(D)** Immunohistochemical analysis of WT and AN iPSC-derived optic cups showing positive staining of PAX6 (green), as well as markers for optic cup domains: VSX2 indicating the neural retina (red, upper panel) and MITF indicating retinal pigmented epithelium (RPE) (red, lower panel). DAPI staining (blue) shows cell nuclei. Scale bar 100μm**. (E)** PAX6 protein analysis detected by western blot in WT and AN iPSC-OCs from day 20 to 35 of differentiation (5-day intervals). PAX6/ACTB ratio was normalised to WT. n=3 (** p<0.01, t-test analysis).

### AN iPSC-derived optic cups show reduced PAX6 protein levels

AN-iPSCs as well as two independent iPSC lines derived from unaffected healthy controls (WT) were further differentiated into 3D optic cup-like stage by adapting established protocols ^29,30^ (Figure 1B). Differentiating organoids showed upregulation of eye field transcription factors (EFTFs) *RAX* and *PAX6* from day 10 onwards (Figure 1C). Organoids were grown until day 35, the timepoint where they mimic an optic cup (OC) like stage, when both neural retina marker VSX2 (Visual system homeobox 2) and retinal pigmented epithelium (RPE) marker MITF (Microphthalmia-associated transcription factor) are present (Figure 1D) ^31,32^. No significant differences in *PAX6* mRNA levels were detected between AN- and WT-iPSC-OCs throughout the process, although a downregulation compared to WT is apparent from day 15 (Figure 1C). Nonetheless, protein immunoblotting showed significant reduction of PAX6 protein in AN iPSC-OCs at day 35, with approx. 0.33 ± 0.23 fold of WT levels (p<0.01, Figure 1E).

### Establishment and characterization of AN iPSCs-derived limbal epithelial stem cells

To test the clinical potential of TRIDs as possible therapy for aniridia limbal stem cell deficiency, the AN iPSC line was further differentiated into 2D limbal epithelial stem cells (LESCs) ^33^. Following formation of embryoid bodies (EBs), limbal fate was induced for 5 days and EBs were plated onto collagen IV coated pates, where epithelial-like cells emerged and proliferated until day 15 (Figure 2A). Timepoint analysis confirmed high expression of LESCs specific markers *ΔNP63α, KRT14* and *ABCG2* by day 15 in AN and WT control lines, proving limbal commitment (Figure 2B). Similar to our 3D optic cup model, no clear differences in *PAX6* mRNA expression were detected between our AN and WT lines (Figure 2C); however, a reduction in full length PAX6 was seen in AN iPSC-LESCs after protein analysis at day 10 and particularly at day 15 (0.37 ± 0.19 fold vs WT=1, p<0.05) (Figure 2D).

**Figure 2.**
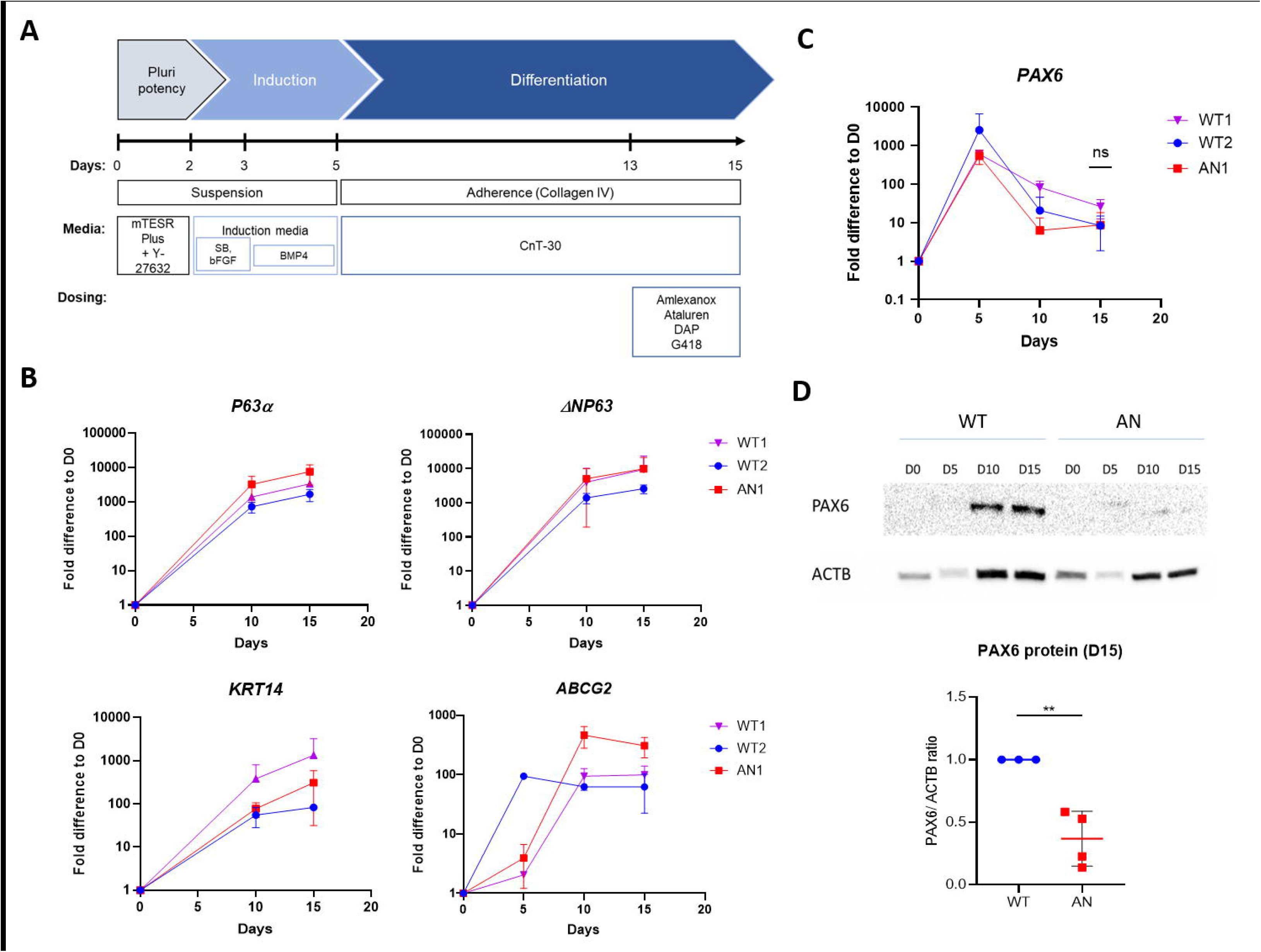
Figure 2. Characterisation of iPSC-derived limbal epithelial stem cell (LESC) from aniridia patient. **(A)** Schematic representation of differentiation protocol used in this study, based on ^33^. **(B)** qRT-PCR transcript analysis of LESC markers *ΔNP63a* (measured with 2 primer pairs), *KRT14* and *ABCG2* in aniridia (AN) and 2 independent control (WT1 and WT2) iPSC lines showing limbal commitment by day 15 of differentiation. **(C)** qRT-PCR transcript analysis of *PAX6* showed no significant difference in expression in AN vs WT lines. Values were normalised to day 0 and to internal housekeeper gene *GAPDH*. Data represent means and SD of n=3 biological replicates. **(D)** Protein analysis detected by western blot revealed decreased PAX6 levels between AN and WT samples from day 10, being statistical significant on day 15 of differentiation. PAX6/ACTB ratio was normalised to control (WT). n=3 (* p<0.05, t-test analysis).

### Amlexanox and Ataluren increase full-length PAX6 levels in aniridia iPSC-optic cups

To test the potential of TRIDs to increase full length PAX6 levels, AN iPSC-OCs were dosed with readthrough compounds amlexanox and DAP, as well as ataluren and G418 from day 15 until collection on day 35 (Figure 1B). G418 caused cell toxicity, even when lower concentrations were tested; the same scenario was observed after DAP dosing, with no viable cells found after day 20 (not shown). The same occurred when dosing WT iPSC-OCs with both drugs, pointing towards drug-specific toxicity. In contrast, amlexanox and ataluren were well tolerated, and no major morphological differences in optic-cup structures were found after dosing (Figure 3A).

**Figure 3.**
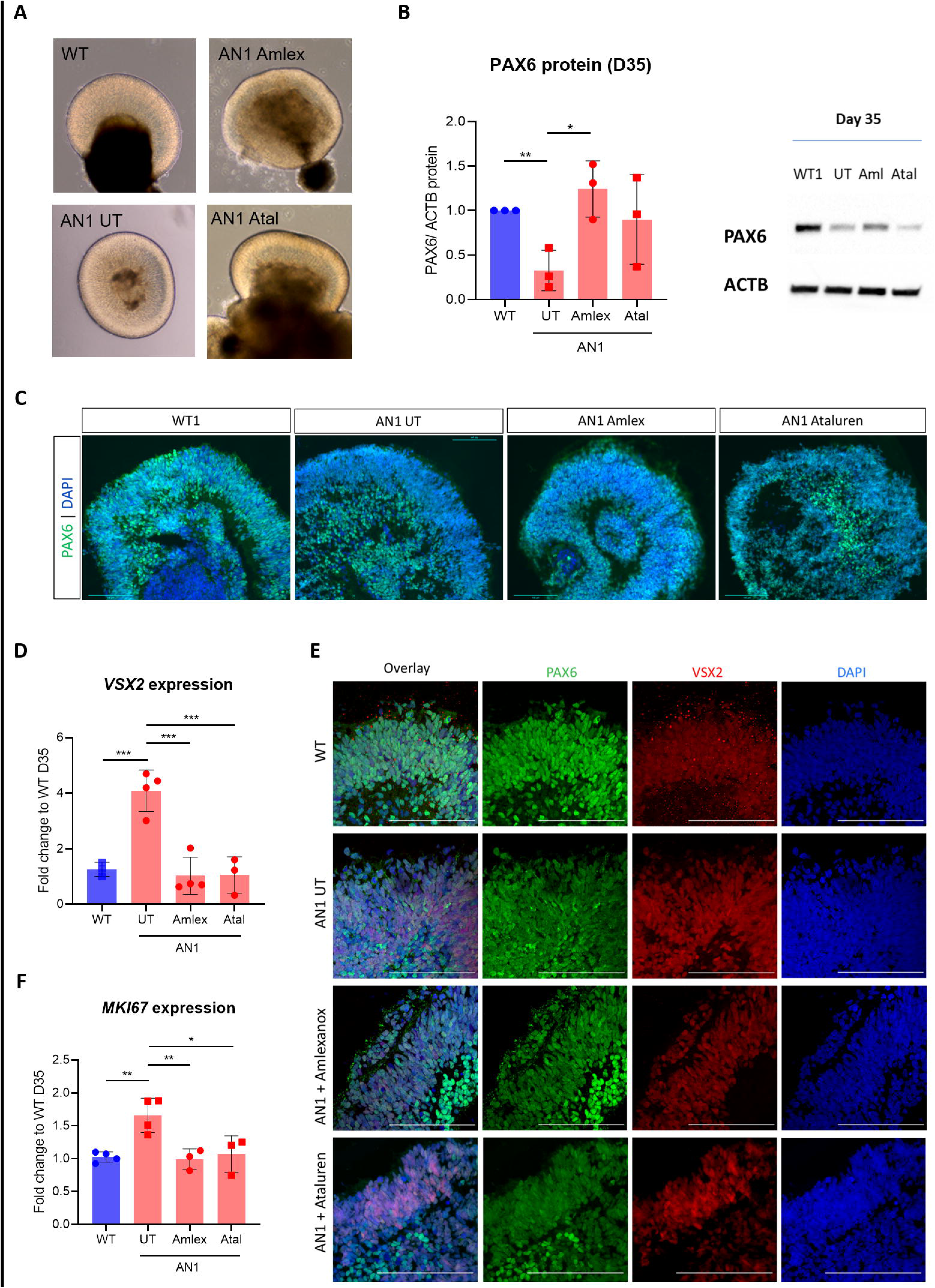
Effect of Translational readthrough inducing drugs (TRIDs) in day 35 aniridia iPSC-derived optic cups (iPSC-OCs). **(A)** Bright-field images of control (WT), untreated aniridia (AN UT), amlexanox-treated aniridia (AN Amlex) and ataluren-treated aniridia (AN Atal) iPSC-OCs. Scale bar 100μm. **(B)** Quantification of PAX6 protein in treated vs untreated AN iPSC-OCs (red bars). PAX6/ACTB ratio was normalised to control (WT, blue bar). (*, p<0.05; **, p<0.01, one-way ANOVA). Data represent means and SD of at least n=3 biological replicates. **(C)** Immunofluorescence analysis on day 35 of differentiation, showing PAX6 staining (green) in control (WT), untreated aniridia (AN UT), amlexanox-treated aniridia (AN Amlex) and ataluren-treated aniridia (AN Atal) iPSC-OCs. DAPI staining is in blue. Scale bar 100μm. **(D)** qRT-PCR transcript analysis of neural retina marker *VSX2* in WT (blue bar) and AN UT, AN Amlex and AN Atal samples (red bars). (*, p<0.05; **, p<0.01, one-way ANOVA). **(E)** Immunohistochemical analysis showing VSX2 staining in WT, AN UT, amlexanox-and ataluren-treated iPSC-OCs. DAPI staining in blue, Scale bar 100μm. **(F)** qRT-PCR transcript analysis of proliferation marker *MKi67* in WT (blue bar) and AN UT, AN Amlex and AN Atal samples (red bars). (***, p<0.001, one-way ANOVA). **(D, F)** Values were normalised to WT and to internal housekeeper gene *GAPDH*. Data represent means and SD of at least n=3 biological replicates.

Full length PAX6 was detected by Western blot in WT, dosed and undosed AN iPSC-OC samples on day 35. We observed that 250μM amlexanox treatment increased full length PAX6 levels by nearly 4-fold (1.24 ± 0.31, p<0.05) compared to untreated AN samples. There was a relative increase in PAX6 in ataluren-treated samples, but it did not reach statistical significance (0.90 ± 0.50, p= 0.22) (Figure 3B). Immunostaining confirmed this result, with untreated AN showing weaker PAX6 staining in the neural retina layer of untreated AN organoids, which improved after treatment with amlexanox (Figure 3C).

### Phenotype rescue in TRID-treated aniridia iPSC-OCs

In order to determine if the increased protein levels following treatment with amlexanox resulted in a functional PAX6 rescue as well as improvement in molecular and cellular phenotype, we investigated the expression of key optic cup marker VSX2 In vivo, VSX2 is necessary for the establishment of retinal progenitor cells (RPCs) in the optic cup and, in the total absence of PAX6, VSX2 expression, along with optic vesicle progression into the optic cup, is abrogated ^13^. Interestingly, the AN iPSC-OCs showed a 4.08 ± 0.74 fold increase of *VSX2* mRNA levels (p<0.001) and immunostaining confirmed a stronger VSX2 signal in untreated AN compared to WT iPSC-OCs (Figure 3D,E). After both amlexanox and ataluren treatment, *VSX2* expression was significantly downregulated to 1.02 ± 0.67 (p<0.001) and 1.05 ± 0.65 -fold (p<0.001), respectively, which was indistinguishable from the levels detected in WT samples (WT expression =1) (Figure 3D). Similarly, immunostaining on day 35 showed weaker VSX2 staining in amlexanox versus untreated AN iPSC-OCs. This was less clear for ataluren-treated AN iPSC-OCs (Figure 3E).

Cell proliferation alterations have been previously reported in response to abnormal *Pax6* levels ^34,35^. Indeed, we observed a significant upregulation in *MKI67* expression, which encodes the proliferation marker Ki-67, in AN iPSC-OCs compared to WT (1.65 ± 0.26 fold, p<0.01). This increased proliferative status was also fully rescued after dosing with amlexanox (0.99 ± 0.16, p<0.01) and ataluren (1.07±0.28, p<0.05) (Figure 3F).

### TRIDs increase PAX6 protein and improve phenotype in iPSC-LESCs

Due to reduced PAX6 protein levels already detected in AN iPSC-LESCs at day 15, we dosed cells for 48h, from day 13 until harvest on day 15 ^23^. Cells treated with 250 μM amlexanox showed affected viability, so lower concentrations, 100 μM and 200 μM, were used. DAP concentrations of 100 μM and 200 μM did not affect cell viability, neither did 40 μM ataluren. In contrast, G418 caused significant cell death in iPSC-LESCs, even at doses lower than 100μg/mL, hence readthrough effect could not be analysed. This was similar to that observed in the 3D optic cup models, confirming G418 cytotoxicity ^9,36^. Overall, TRIDs dosing increased full-length PAX6 in AN iPSC-LESCs (Figure 4A): amlexanox significantly improved protein levels to 0.650 ± 0.043 - fold (100 μM, p<0.05) and 0.941 ± 0.085-fold (200 μM, p<0.0001). Also 100 μM DAP treatment improved PAX6 levels to 0.912 ± 0.064 (p<0.001). Ataluren treated cells also showed significant increase in PAX6, with full-length levels reaching 0.85±0.048-fold (p<0.001) of control levels (Figure 4A).

**Figure 4.**
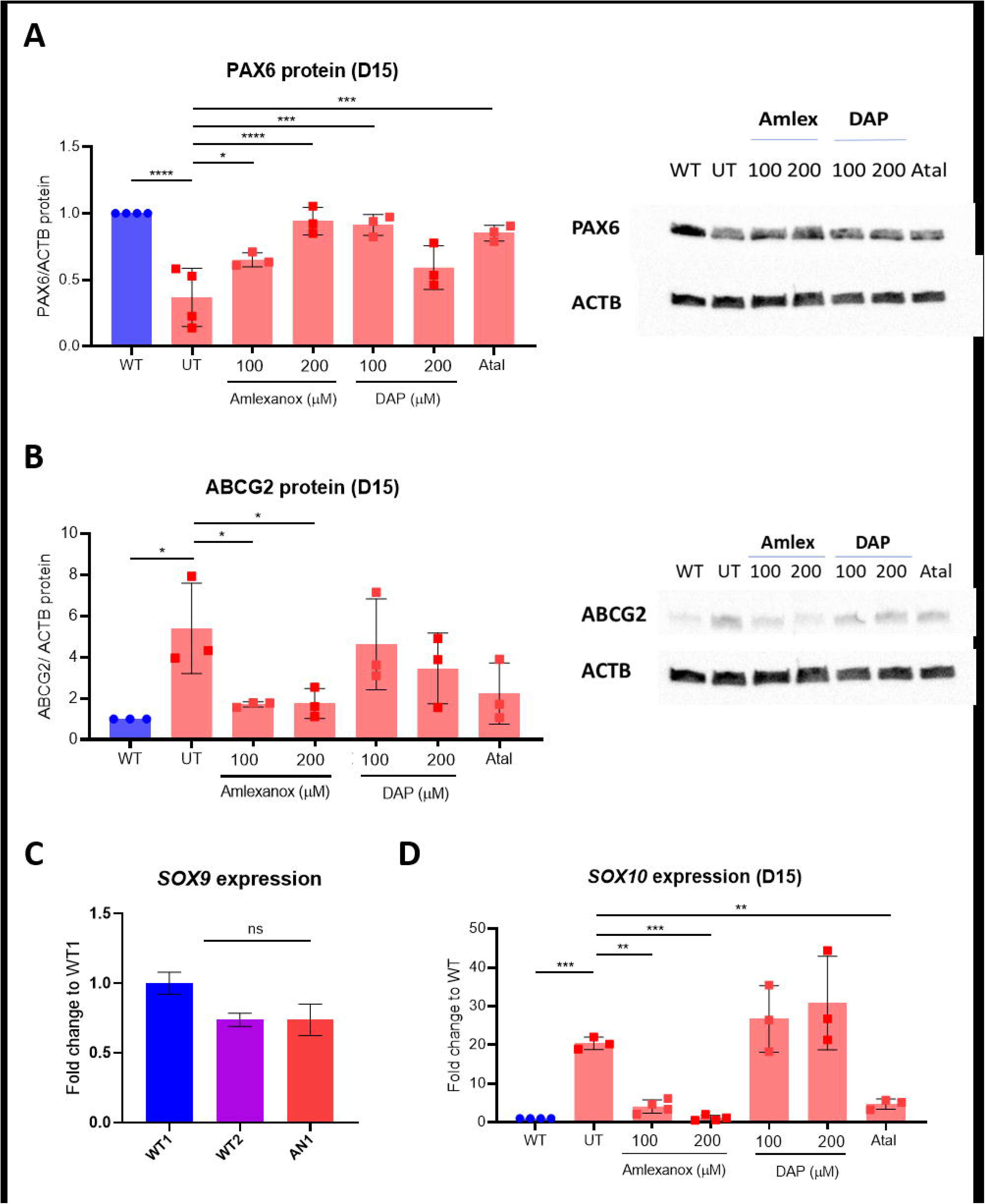
TRIDs rescue PAX6 expression in aniridia iPSC-derived LESCs. **(A)** Quantification of PAX6 protein in untreated (UT) AN iPSC-LESCs versus amlexanox-, DAP- and ataluren-treated AN iPSC-LESCs (red bars). PAX6/ACTB ratio was normalised to control (WT, blue bar). Values in x axis refer to compounds concentrations in μM. (*, p<0.05; ***, p<0.001; ****, p<0.0001; one-way ANOVA). Data represent means and SD of at least n=3 biological replicates. (B) Quantification of ABCG2 protein detected by western blot in WT (blue bar), AN untreated and amlexanox-, DAP- and ataluren-treated AN iPSC-LESCs (red bars). Values in x axis refer to compounds concentrations in μM. PAX6/ACTB ratio was normalised to control (WT). Data represent means and SD of n=3 biological replicates (*, p<0.05, one-way ANOVA). **(C)** Relative expression of *SOX9* transcripts in WT1 and WT2 and AN iPSC-LESCs. Significance was calculated using multiple t-test between AN and both WT lines. **(D)** qRT-PCR transcript analysis of neural crest marker *SOX10* in WT (blue bar) and AN UT, AN Amlex and AN Atal samples (red bars). (***, p<0.001, **, p<0.01; one-way ANOVA). **(C, D)** Values were normalised to WT and to internal housekeeper gene *GAPDH*. Data represent means and SD of at least n=3 biological replicates.

To assess for functional and phenotypic rescue following treatment with TRIDs we examined expression of ABCG2, which is transiently expressed in LESCs and is considered a LESC-specific stemness marker ^37^. Although the relationship between PAX6 and ABCG2 is not known, it was recently shown that *ABCG2* mRNA is upregulated in LESCs extracted from aniridia patients (with PTC-causing mutations) compared to controls ^38^. We observed similar results in our iPSC-derived system, where *ABCG2* mRNA peaked at day 10 in all lines (Figure 3B) and, at day 15, there was a 5.40 ±1.79 fold (p<0.01) accumulation of ABCG2 protein in AN compared to WT control iPSC-LESCs (p<0.01) (Figure 4B). Remarkably, amlexanox-treated AN iPSC-LESCs showed a very significant reduction in ABCG2 protein, reaching levels very close to WT, with both concentrations: 100μM (1.70 ± 0.10 -fold, p<0.05) and 200μM (1.75 ± 0.59, p<0.05) (Figure 4B).

SOX9 and SOX10 are TFs expressed in the neural crest-fate cells in the limbal niche, which is essential for the homeostasis of LESCs ^16^. Therefore, and because PAX6 was recently shown to drive neural crest migration during corneal development ^15,17^, we tested the expression of *SOX9* and *SOX10* between AN and WT iPSC-LESCs. Although *SOX9* expression was not significantly altered between AN and WT (p= 0.353, Figure 4C), we found that *SOX10* was sharply upregulated in AN iPSC-LESCs (20 ±1.29 -fold, p<0.001, Figure 4D). Following treatment with TRIDs, *SOX10* expression was rescued by 100μM (4.06± 1.56, p<0.01) and 200μM (1.07 ± 0.60, p=0.001) of amlexanox, as well as with ataluren (6.26±2.88, p<0.01) (Figure 4D). Similarly to previous results, DAP did not induce an improvement in *SOX10* expression (Figure 4D).

In conclusion, amlexanox increases full-length PAX6 levels and rescues phenotypic differences in both early 3D optic cups and 2D limbal epithelial stem cells generated from aniridia patient iPSCs, proving that newly synthesized PAX6 is functional and the new amino acid inserted is likely tolerated.

## Discussion

The aim of this work is to provide proof of principle for the further development of repurposed readthrough drugs amlexanox and 2,6-diaminopurine (DAP) for aniridia. Aniridia is a highly suitable disease for readthrough therapy approaches, due to the high prevalence of *PAX6* nonsense mutations, dosage sensitivity, and if the target tissue is well considered i.e. cornea and LESC to reduce aniridia-related keratopathy (ARK) and maintain levels of vision. Insufficient PAX6 levels, or haploinsufficiency, is thought to be the underlying genetic mechanism of aniridia; therefore, increasing full-length PAX6 levels, even if not fully, might be enough to attenuate disease. This is also supported in patients, where PTC-introducing variants are generally associated with severe forms of aniridia, whilst patients with missense mutations usually present with milder phenotypes and less severe vision loss ^2,3,5^.

We generated an iPSC line from an aniridia patient carrying the heterozygous *PAX6* nonsense mutation c.781C>T, p.(Arg261*). This variant is located within the “PAX6 mutation hotspot”, a region in exons 8 to 13 with methylated CpG islands, where 21% of all mutations and 60% of all nonsense mutations are located ^8,39,40^.

We have differentiated the patient iPSCs into 3D optic cups (iPSC-OCs) and show significantly reduced PAX6 protein levels at day 35, a timepoint comparable to the *in vivo* optic cup stage. Nonetheless, these reduced PAX6 levels are sufficient to form optic cup domains (neural retina and RPE) in our *in vitro* system, which is also consistent with *in vivo* results ^13^. Amlexanox and ataluren were shown to recover levels of full-length PAX6, while DAP and G418 showed toxicity at all concentrations tested.

We observed a striking increase in neural retina marker VSX2 expression in aniridia iPSC-derived optic cups. Low Pax6 levels seem to promote early neurogenesis in the mice optic vesicle ^13^; this might explain the accumulation of VSX2, which was detected at both mRNA and protein levels. Importantly, we observe that normal VSX2 levels are restored after treatment of aniridia iPSC-OCs with amlexanox and ataluren. These results suggest that both compounds induce functional PAX6 protein increase, leading to rescue of the *in vitro* phenotype. This was further supported by the downregulation of proliferation marker *MKi67* expression, known to be increased in *Pax6* mutant cells, after dosing of aniridia iPSC-OCs with both TRIDs ^34^.

The role of PAX6 in the eye is both time- and tissue-specific, acting during development but also on maintenance of adult tissue ^13,41^. This translates into a developmental and progressive disease, where aniridia patients typically show hypoplasia of the iris and fovea from birth, and progressive opacity of the lens and cornea from childhood/early adulthood ^1,8^. From large natural history studies we understand that the visual acuity remains relatively stable over decades of life ^3^. Therapeutic approaches targeting developmental defects are currently not feasible, hence we aimed to test the clinical potential of TRIDs to halt or slow down ARK, which can affect up to 90% patients and is the mainstay for a decline in visual acuity over time ^8^. For that purpose, we established a second aniridia human model, by growing patient iPSC-derived 2D limbal epithelial stem cells (iPSC-LESCs). Upregulation of LESCs specific markers *ΔNP63α, ABCG2* and *KRT14* in these cells proved commitment to the limbal fate. Aniridia patient iPSC-LESCs show an approx. 60% reduction in PAX6 protein levels, lower than the estimated 50%, validating this model to study *PAX6* haploinsufficiency. Once again, we did not observe significantly reduced *PAX6* transcript levels in AN vs WT iPSC-LESCs during the first 15 days of differentiation. It is assumed that *PAX6* null variants lead to the degradation of the mutated transcripts via NMD, thus resulting in haploinsufficiency ^8^; however, we do not seem to observe this in our in vitro models.

Dosing of AN iPSC-LESCs with different TRIDs resulted in similar profiles compared to 3D optic cups; both amlexanox and ataluren proved to significantly increase PAX6 protein, although slightly lower concentrations of amlexanox were used due to very low proliferation in cells treated with 250μM. Importantly, both compounds, but particularly amlexanox, induced strong phenotype rescue by restoring ABCG2 as well as *SOX10* levels, two important players in limbal epithelial stem cell identity and survival, respectively ^16,37^. Although DAP also induced a significant increase in PAX6 levels in AN iPSC-LESCs, it did not show a significant downstream phenotypic rescue. We hypothesize this is due to the new amino acid introduced, i.e. tryptophan, which may still have a deleterious effect ^26^. Ataluren was well tolerated in both models; in contrast, G418 was highly cytotoxic, proving the downside of traditional aminoglycosides use and need for less toxic TRIDs ^9^.

ABCG2 is a transient limbal epithelial stem cell marker, turned off when cells exit the stem cell state and start differentiation into corneal epithelial cells ^37^. We hypothesize that ABCG2 increased levels in AN iPSC-LESCs show that these cells may be unable to either switch off their proliferative status and/or trigger the differentiation process into corneal epithelial cells ^38^. In parallel, we observe altered expression of neural crest marker *SOX10*, supporting the recent evidence that PAX6 has a role in neural crest-derived cells from the limbal niche ^15,17^. Further differentiation of AN iPSC-LESCs into later stages as well as high throughput molecular characterisation of these cells would be important to not only understand the mechanisms behind PAX6-related LSCD but also to understand how iPSC-derived models compare to *in vivo* development and disease, particularly when dealing with such a regulatory-complex transcription factor like PAX6.

Overall, patients with missense *PAX6* mutations tend to have milder ocular phenotypes ^2,5^; in our recently published 86 aniridia patient cohort, patients with missense mutations have significantly lower incidence of ARK, compared to patients with nonsense variants, who present with the highest ARK prevalence ^3^. However, the nearly complete absence of aniridia patients with missense mutations located downstream of exon 7, coupled with the variable expressivity of the disease, makes it difficult to accurately predict the genotype-phenotype relationships. The closest reported missense variant to c.781C>T, p.(Arg261*) located in exon 10 (predicted homeodomain) was c.773T>C, p.(Phe258Ser); the patient presented with typical iris hypoplasia, and chorioretinal coloboma involving the optic disc, but indeed no description of corneal problems ^42^.

In this study we provide strong evidence supporting the repurposing of amlexanox as a putative therapeutic compound for aniridia patients with *PAX6* nonsense mutations. Our 3D optic cup models showed good tolerance to amlexanox, but in order to reduce off-target effects or systemic complications, topical formulations with a lower dose could be administered ^43^. Further work on higher order *in vivo* models may be required to ascertain the optimal dose needed to induce optimal readthrough in aniridia patients. Interestingly, amlexanox has recently been shown to improve glucose levels and enhance liver fat loss in individuals with type II diabetes ^44^. Hence, it could be beneficial to assess the effect of systemic amlexanox in aniridia patients, since recent reports show that aniridia patients commonly present with metabolic dysregulation leading to obesity and type II diabetes ^3,45^. Therefore, we speculate that amlexanox might have ocular and wider systemic benefit in aniridia patients.

Lastly, this work provides further evidence that readthrough therapy seems to be a particularly promising therapeutic approach for aniridia, with previous *in vivo* models ^11,12^ and now patient-specific *in vitro* models showing positive pre-clinical outcomes. The advances in readthrough drug development allied to more complex and representative human disease models will certainly allow for new compounds to be pushed into clinical trials for aniridia patients.

## Materials and Methods

### Ethics and clinical description

This study was approved by Moorfields Eye Hospital and the National Research Ethics Committee and was conducted in adherence to the tenets of the Declaration of Helsinki; informed written consent was obtained from all participants. A 4-mm punch skin biopsy was obtained from a 6-year-old male aniridia patient (genotype confirmed *PAX6* c.781C>T, p.(Arg261*) from the upper arm. The patient was hypermetropic (right eye +6.00/-2.00×10 and left eye +6.00/−1.75×180) and their best corrected visual acuity was 0.74 LogMAR in each eye. Intraocular pressure was within normal range (18 mmHg in both eyes), no signs of glaucoma, cataracts or ARK, both cornea were clear. The patient does have complete iris and foveal hypoplasia.

### Induced Pluripotent Stem Cells (iPSC) generation

AN patient iPSCs were generated using non-integrating episomal reprogramming of dermal fibroblasts extracted from a skin biopsy from the patient’s arm, following established protocols ^27,46^. A minimum of 2 clonal lines were expanded and characterised as previously described ^28,46^. Control (WT) iPSCs used in this study were previously published ^46^.

All iPSC lines were maintained in mTESR Plus media (StemCell Technologies, Canada) with 0.1% Pen/Strep on Matrigel-coated wells (1:100) (Corning, USA). For passaging, ReLESR (StemCell Technologies, Canada) was used for detaching and after 24h, iPSCs were fed daily with mTESR Plus until confluent.

### iPSC-differentiation into 3D optic cups

Differentiation of iPSCs into 3D optic cups was performed based on published protocols ^29,30^. Briefly, confluent iPSCs were detached with Accumax (ThermoFisher Scientific, USA) to single cell suspension and 3.6 million cells per well were plated onto Aggrewell400 plates (StemCell Technologies, Canada) (3,000 cells per microwell) in mTESR Plus with 10μM Y-27632 (Abcam), following manufacturer’s instructions. After 48h, embryoid bodies (EBs) were collected and plated onto low attachment 60mm plates in neural induction media (NIM) – DMEM/F12 (ThermoFisher Scientific), 20% knock-out serum replacement (KOSR) (ThermoFisher Scientific), 2% B27 (ThermoFisher Scientific), 1x Non-essential amino acids (NEAA; ThermoFisher Scientific), 1% Pen/Strep, 1xGlutamax (ThermoFisher Scientific) and 5ng/mL IGF-1 (Sigma-Aldrich) – until day 7. On day 8, cells were cultured in NIM with 15% KOSR and finally with 10% KOSR from day 11 until day 35 (Figure 1B).

### iPSC differentiation into LESCs

Differentiation of iPSCs into LESCs was done following the protocol from Hongisto et al, with small adjustments ^33^. Confluent iPSCs (~90/95%) were detached using ReLESR and clumps resuspended in mTESR Plus with 10μM Y-27632. Cell clumps were transferred into non-coated (petri) dishes and incubated O/N to allow the formation of EBs (Day 0). After 48h (Day 2), EBs were carefully washed with DPBS and resuspended in SM media (KnockOut DMEM supplemented with 15% xeno-free serum replacement, 2mM L-glutamine, 0.1mM 2-mercaptoethanol, 1% non-essential amino acids, and 50U/mL penicillin-streptomycin) supplemented with 10μM SB-505124 (Sigma Aldrich, USA) and 50ng/mL bFGF (Peprotech, USA). Media was replaced with SM media supplemented with 25ng/mL BMP-4 (Peprotech, USA) on days 3 and 4. On day 5, EBs were carefully plated into collagen IV-coated wells in a mix of CnT30 media (CellnTech, Switzerland) and SM media (3:1) and allowed to attach for 48h. From there on, media was changed with CnT30 every other day until collection on day 15 for RNA and protein analysis.

### Dosing and compounds information

Dosing concentrations of amlexanox (Abcam) and 2,6-diaminopurine (DAP, Sigma-Aldrich) were based on previous publications ^23,25,26^. Known TRIDs ataluren/ PTC124 (ApexBio Tech LLC) and G418 (Life Technologies) were used as positive readthrough controls at 40μM and 100μg/mL, respectively, according to previous publications from our group ^24,47,48^. 3D optic cups were dosed with amlexanox 250μM or ataluren 40μM in NIM+10%KOSR from day 15 to 35 of differentiation, with media refreshed every other day. iPSC-LESCs were dosed from day 13 to 15 of differentiation in CnT-30 media, with media change after 24h. Amlexanox (100μM and 200μM), DAP (100μM and 200μM), Ataluren (40μM) and G418 (100μg/mL) were tested.

### RNA extraction and RT-qPCR

For transcript analysis of 3D optic cups, RNA extraction was performed after pellets collection using the RNeasy Mini or Micro Kit (QIAGEN, Germany); iPSC-LESCs were harvested by adding 300μL of Lysis buffer (Zymo Research, USA) and cells collected using a cell scraper; RNA was extracted following instructions in the Quick-RNA™ MicroPrep Kit w/ Zymo-Spin™ IC Columns kit (Zymo Research).

cDNA was synthesized from 500ng RNA using High-Capacity RNA-to-cDNA Kit (Life Technologies). RT-qPCR was performed with 2x SYBR Green MasterMix (ThermoFisher Scientific, USA) on a StepOne Real-Time PCR system (Applied Biosystems, UK) or QuantStudio 6 Flex (Applied Biosystems, UK). Primers used for qRT-PCR are listed in Table S1. Transcript levels were measured in duplicate and normalised to housekeeper genes *GAPDH or ACTB*. The relative expression of each target gene was calculated using the comparative C_T_ method.

### Western blotting

Samples were analysed by western blotting as described previously ^24,49^. Cells were washed with ice-cold PBS and total protein extract was prepared with RIPA buffer with 1x Halt ™ protease inhibitor cocktail and Halt ™ phosphatase inhibitor (ThermoFisher Scientific, MA, USA) at a ratio of 5×106 cells/mL. 30μg protein for iPSC-derived optic cups or 15μg for iPSC-LESCs were loaded onto 4-15% Mini-PROTEAN ^®^ TGX ™ gels (BioRad Inc., CA, USA) and transferred to an Immun-Blot™ PVDF membrane using a Trans-Blot^®^ SD semi-dry transfer cell (BioRad Inc., CA, USA). Membranes were blocked with 5% non-fat dry milk in PBST for 2h, incubated overnight at 4°C with the following primary antibodies diluted in blocking buffer: PAX6 (1:2000, Covance); ABCG2 (1:1000, SantaCruz); ACTB (1:5000, SigmaAldrich). Incubation with horseradish peroxidase conjugated secondary antibody anti-mouse or rabbit 1:5000 (Applied Biosystems, UK) was done for 2h at room temperature. Membranes were incubated with Clarity Western ECL Substrate (BioRad Inc., CA, USA) and imaged using the ChemiDoc XRS™ Imaging System (BioRad Inc., CA, USA). Band intensities were quantified using the Fiji/ImageJ software (National Institutes of Health, MD, USA).

### Immunofluorescence and Imaging

Day 35 iPSC-OCs were processed for immunohistochemistry analysis following the protocol from Reichmann et al ^50^. Slides were imaged using an EVOS FL system (ThermoFisher Scientific, USA) and ZEISS LSM 700 or LSM 710 (ZEISS Research, Germany).

### Statistical Analysis

Statistical analysis was performed using GraphPad Prism 8.0 (GraphPad Software Inc., San Diego, CA, USA). One-Way ANOVA with multiple comparisons was used for comparison studies, with significance achieved with *p* value of ≤ 0.05 (*), ≤ 0.01 (**), ≤ 0.001 (***). All results are expressed as mean ± SD, unless specified. Experiments were performed with n = 3 biological replicates, except when specified.

## Acknowledgments

The authors would like to thank the patients and their families for donating skin samples. Funding for this work was obtained from Moorfields Eye Charity, Wellcome Trust (Grant 205174/Z/16/Z), Fight for Sight and Aniridia Network to MM; NWO Visitor Travel Grant (Grant nr 040.11.699) to JZ and DLC.

## Author Contributions

MM designed research and acquired funding; DLC, PH, JH performed research; JZ contributed with reagents and research design; DLC and MM analysed data; DLC wrote the first draft; DLC and MM edited the manuscript with input from all authors.

## Data Availability Statement

Data sharing not applicable to this article as no datasets were generated or analysed during the current study.

## References

1. Moosajee, M., Hingorani, M., and Moore, A.T. (2018). PAX6-Related Aniridia. In GeneReviews(®), (Seattle (WA): University of Washington, Seattle).

2. Hingorani, M., Williamson, K.A., Moore, A.T., and van Heyningen, V. (2009). Detailed ophthalmologic evaluation of 43 individuals with PAX6 mutations. Investigative ophthalmology & visual science 50, 2581–2590. 10.1167/iovs.08-2827.

3. Kit, V., Lima Cunha, D., Hagag, A.M., and Moosajee, M. (2021). Longitudinal genotype-phenotype analysis in 86 PAX6-related aniridia patients. JCI insight. 10.1172/jci.insight.148406.

4. Latta, L., Figueiredo, F.C., Ashery-Padan, R., Collinson, J.M., Daniels, J., Ferrari, S., Szentmáry, N., Solá, S., Shalom-Feuerstein, R., Lako, M., et al. (2021). Pathophysiology of aniridia-associated keratopathy: Developmental aspects and unanswered questions. The ocular surface 22, 245–266. 10.1016/j.jtos.2021.09.001.

5. Lagali, N., Wowra, B., Fries, F.N., Latta, L., Moslemani, K., Utheim, T.P., Wylegala, E., Seitz, B., and Käsmann-Kellner, B. (2020). PAX6 Mutational Status Determines Aniridia-Associated Keratopathy Phenotype. Ophthalmology 127, 273–275. 10.1016/j.ophtha.2019.09.034.

6. Jordan, T., Hanson, I., Zaletayev, D., Hodgson, S., Prosser, J., Seawright, A., Hastie, N., and van Heyningen, V. (1992). The human PAX6 gene is mutated in two patients with aniridia. Nature genetics 1, 328–332. 10.1038/ng0892-328.

7. Hill, R.E., Favor, J., Hogan, B.L., Ton, C.C., Saunders, G.F., Hanson, I.M., Prosser, J., Jordan, T., Hastie, N.D., and van Heyningen, V. (1991). Mouse small eye results from mutations in a paired-like homeobox-containing gene. Nature 354, 522–525. 10.1038/354522a0.

8. Lima Cunha, D., Arno, G., Corton, M., and Moosajee, M. (2019). The Spectrum of PAX6 Mutations and Genotype-Phenotype Correlations in the Eye. Genes 10. 10.3390/genes10121050.

9. Way, C.M., Lima Cunha, D., and Moosajee, M. (2020). Translational readthrough inducing drugs for the treatment of inherited retinal dystrophies. Expert Review of Ophthalmology 15, 169–182. 10.1080/17469899.2020.1762489.

10. Richardson, R., Smart, M., Tracey-White, D., Webster, A.R., and Moosajee, M. (2017). Mechanism and evidence of nonsense suppression therapy for genetic eye disorders. Experimental eye research 155, 24–37. 10.1016/j.exer.2017.01.001.

11. Gregory-Evans, C.Y., Wang, X., Wasan, K.M., Zhao, J., Metcalfe, A.L., and Gregory-Evans, K. (2014). Postnatal manipulation of Pax6 dosage reverses congenital tissue malformation defects. The Journal of clinical investigation 124, 111–116. 10.1172/jci70462.

12. Wang, X., Gregory-Evans, K., Wasan, K.M., Sivak, O., Shan, X., and Gregory-Evans, C.Y. (2017). Efficacy of Postnatal In Vivo Nonsense Suppression Therapy in a Pax6 Mouse Model of Aniridia. Molecular therapy. Nucleic acids 7, 417–428. 10.1016/j.omtn.2017.05.002.

13. Shaham, O., Menuchin, Y., Farhy, C., and Ashery-Padan, R. (2012). Pax6: a multi-level regulator of ocular development. Progress in retinal and eye research 31, 351–376. 10.1016/j.preteyeres.2012.04.002.

14. Collinson, J.M., Chanas, S.A., Hill, R.E., and West, J.D. (2004). Corneal development, limbal stem cell function, and corneal epithelial cell migration in the Pax6(+/−) mouse. Investigative ophthalmology & visual science 45, 1101–1108. 10.1167/iovs.03-1118.

15. Chen, S.Y., Cheng, A.M.S., Zhang, Y., Zhu, Y.T., He, H., Mahabole, M., and Tseng, S.C.G. (2019). Pax 6 Controls Neural Crest Potential of Limbal Niche Cells to Support Self-Renewal of Limbal Epithelial Stem Cells. Scientific reports 9, 9763. 10.1038/s41598-019-45100-7.

16. Su, Z., Wang, J., Lai, Q., Zhao, H., and Hou, L. (2020). KIT ligand produced by limbal niche cells under control of SOX10 maintains limbal epithelial stem cell survival by activating the KIT/AKT signalling pathway. Journal of cellular and molecular medicine 24, 12020–12031. 10.1111/jcmm.15830.

17. Takamiya, M., Stegmaier, J., Kobitski, A.Y., Schott, B., Weger, B.D., Margariti, D., Cereceda Delgado, A.R., Gourain, V., Scherr, T., Yang, L., et al. (2020). Pax6 organizes the anterior eye segment by guiding two distinct neural crest waves. PLoS genetics 16, e1008774. 10.1371/journal.pgen.1008774.

18. Takahashi, K., Tanabe, K., Ohnuki, M., Narita, M., Ichisaka, T., Tomoda, K., and Yamanaka, S. (2007). Induction of pluripotent stem cells from adult human fibroblasts by defined factors. Cell 131, 861–872. 10.1016/j.cell.2007.11.019.

19. Hata, M., Ikeda, H.O., Iwai, S., Iida, Y., Gotoh, N., Asaka, I., Ikeda, K., Isobe, Y., Hori, A., Nakagawa, S., et al. (2018). Reduction of lipid accumulation rescues Bietti’s crystalline dystrophy phenotypes. Proceedings of the National Academy of Sciences of the United States of America 115, 3936–3941. 10.1073/pnas.1717338115.

20. Lane, A., Jovanovic, K., Shortall, C., Ottaviani, D., Panes, A.B., Schwarz, N., Guarascio, R., Hayes, M.J., Palfi, A., Chadderton, N., et al. (2020). Modeling and Rescue of RP2 Retinitis Pigmentosa Using iPSC-Derived Retinal Organoids. Stem cell reports 15, 67–79. 10.1016/j.stemcr.2020.05.007.

21. Ramsden, C.M., Nommiste, B., A, R.L., Carr, A.F., Powner, M.B., M, J.K.S., Chen, L.L., Muthiah, M.N., Webster, A.R., Moore, A.T., et al. (2017). Rescue of the MERTK phagocytic defect in a human iPSC disease model using translational read-through inducing drugs. Scientific reports 7, 51. 10.1038/s41598-017-00142-7.

22. Greer, R.O., Jr., Lindenmuth, J.E., Juarez, T., and Khandwala, A. (1993). A double-blind study of topically applied 5% amlexanox in the treatment of aphthous ulcers. Journal of oral and maxillofacial surgery: official journal of the American Association of Oral and Maxillofacial Surgeons 51, 243–248; discussion 248-249. 10.1016/s0278-2391(10)80164-8.

23. Atanasova, V.S., Jiang, Q., Prisco, M., Gruber, C., Piñón Hofbauer, J., Chen, M., Has, C., Bruckner-Tuderman, L., McGrath, J.A., Uitto, J., et al. (2017). Amlexanox Enhances Premature Termination Codon Read-Through in COL7A1 and Expression of Full Length Type VII Collagen: Potential Therapy for Recessive Dystrophic Epidermolysis Bullosa. The Journal of investigative dermatology 137, 1842–1849. 10.1016/j.jid.2017.05.011.

24. Eintracht, J., Forsythe, E., May-Simera, H., and Moosajee, M. (2021). Translational readthrough of ciliopathy genes BBS2 and ALMS1 restores protein, ciliogenesis and function in patient fibroblasts. EBioMedicine 70, 103515. 10.1016/j.ebiom.2021.103515.

25. Gonzalez-Hilarion, S., Beghyn, T., Jia, J., Debreuck, N., Berte, G., Mamchaoui, K., Mouly, V., Gruenert, D.C., Déprez, B., and Lejeune, F. (2012). Rescue of nonsense mutations by amlexanox in human cells. Orphanet journal of rare diseases 7, 58. 10.1186/1750-1172-7-58.

26. Trzaska, C., Amand, S., Bailly, C., Leroy, C., Marchand, V., Duvernois-Berthet, E., Saliou, J.M., Benhabiles, H., Werkmeister, E., Chassat, T., et al. (2020). 2,6-Diaminopurine as a highly potent corrector of UGA nonsense mutations. Nature communications 11, 1509. 10.1038/s41467-020-15140-z.

27. Parfitt, D.A., Lane, A., Ramsden, C.M., Carr, A.F., Munro, P.M., Jovanovic, K., Schwarz, N., Kanuga, N., Muthiah, M.N., Hull, S., et al. (2016). Identification and Correction of Mechanisms Underlying Inherited Blindness in Human iPSC-Derived Optic Cups. Cell stem cell 18, 769–781. 10.1016/j.stem.2016.03.021.

28. Harding, P., Lima Cunha, D., Méjécase, C., Eintracht, J., Toualbi, L., Sarkar, H., and Moosajee, M. (2021). Generation of human iPSC line (UCLi013-A) from a patient with microphthalmia and aniridia, carrying a heterozygous missense mutation c.372C>A p.(Asn124Lys) in PAX6. Stem cell research 51, 102184. 10.1016/j.scr.2021.102184.

29. Mellough, C.B., Collin, J., Khazim, M., White, K., Sernagor, E., Steel, D.H., and Lako, M. (2015). IGF-1 Signaling Plays an Important Role in the Formation of Three-Dimensional Laminated Neural Retina and Other Ocular Structures From Human Embryonic Stem Cells. Stem cells (Dayton, Ohio) 33, 2416–2430. 10.1002/stem.2023.

30. Eintracht, J., Harding, P., Lima Cunha, D., and Moosajee, M. (2022). Efficient embryoid-based method to improve generation of optic vesicles from human induced pluripotent stem cells [version 1; peer review: 1 approved]. 11. 10.12688/f1000research.108829.1.

31. Nakano, T., Ando, S., Takata, N., Kawada, M., Muguruma, K., Sekiguchi, K., Saito, K., Yonemura, S., Eiraku, M., and Sasai, Y. (2012). Self-formation of optic cups and storable stratified neural retina from human ESCs. Cell stem cell 10, 771–785. 10.1016/j.stem.2012.05.009.

32. Zhong, X., Gutierrez, C., Xue, T., Hampton, C., Vergara, M.N., Cao, L.H., Peters, A., Park, T.S., Zambidis, E.T., Meyer, J.S., et al. (2014). Generation of three-dimensional retinal tissue with functional photoreceptors from human iPSCs. Nature communications 5, 4047. 10.1038/ncomms5047.

33. Hongisto, H., Vattulainen, M., Ilmarinen, T., Mikhailova, A., and Skottman, H. (2018). Efficient and Scalable Directed Differentiation of Clinically Compatible Corneal Limbal Epithelial Stem Cells from Human Pluripotent Stem Cells. Journal of visualized experiments: JoVE. 10.3791/58279.

34. Rabiee, B., Anwar, K.N., Shen, X., Putra, I., Liu, M., Jung, R., Afsharkhamseh, N., Rosenblatt, M.I., Fishman, G.A., Liu, X., et al. (2020). Gene dosage manipulation alleviates manifestations of hereditary PAX6 haploinsufficiency in mice. Science translational medicine 12. 10.1126/scitranslmed.aaz4894.

35. Ouyang, J., Shen, Y.C., Yeh, L.K., Li, W., Coyle, B.M., Liu, C.Y., and Fini, M.E. (2006). Pax6 overexpression suppresses cell proliferation and retards the cell cycle in corneal epithelial cells. Investigative ophthalmology & visual science 47, 2397–2407. 10.1167/iovs.05-1083.

36. Hancock, H.A., Guidry, C., Read, R.W., Ready, E.L., and Kraft, T.W. (2005). Acute aminoglycoside retinal toxicity in vivo and in vitro. Investigative ophthalmology & visual science 46, 4804–4808. 10.1167/iovs.05-0604.

37. Vattulainen, M., Ilmarinen, T., Koivusalo, L., Viiri, K., Hongisto, H., and Skottman, H. (2019). Modulation of Wnt/BMP pathways during corneal differentiation of hPSC maintains ABCG2-positive LSC population that demonstrates increased regenerative potential. Stem cell research & therapy 10, 236. 10.1186/s13287-019-1354-2.

38. Latta, L., Nordström, K., Stachon, T., Langenbucher, A., Fries, F.N., Szentmáry, N., Seitz, B., and Käsmann-Kellner, B. (2019). Expression of retinoic acid signaling components ADH7 and ALDH1A1 is reduced in aniridia limbal epithelial cells and a siRNA primary cell based aniridia model. Experimental eye research 179, 8–17. 10.1016/j.exer.2018.10.002.

39. Tzoulaki, I., White, I.M., and Hanson, I.M. (2005). PAX6 mutations: genotype-phenotype correlations. BMC genetics 6, 27. 10.1186/1471-2156-6-27.

40. Wawrocka, A., and Krawczynski, M.R. (2018). The genetics of aniridia - simple things become complicated. Journal of applied genetics 59, 151–159. 10.1007/s13353-017-0426-1.

41. Marquardt, T., Ashery-Padan, R., Andrejewski, N., Scardigli, R., Guillemot, F., and Gruss, P. (2001). Pax6 is required for the multipotent state of retinal progenitor cells. Cell 105, 43–55. 10.1016/s0092-8674(01)00295-1.

42. Azuma, N., Yamaguchi, Y., Handa, H., Tadokoro, K., Asaka, A., Kawase, E., and Yamada, M. (2003). Mutations of the PAX6 gene detected in patients with a variety of optic-nerve malformations. American journal of human genetics 72, 1565–1570. 10.1086/375555.

43. Woodley, D.T., Cogan, J., Hou, Y., Lyu, C., Marinkovich, M.P., Keene, D., and Chen, M. (2017). Gentamicin induces functional type VII collagen in recessive dystrophic epidermolysis bullosa patients. The Journal of clinical investigation 127, 3028–3038. 10.1172/jci92707.

44. Oral, E.A., Reilly, S.M., Gomez, A.V., Meral, R., Butz, L., Ajluni, N., Chenevert, T.L., Korytnaya, E., Neidert, A.H., Hench, R., et al. (2017). Inhibition of IKK⍰ and TBK1 Improves Glucose Control in a Subset of Patients with Type 2 Diabetes. Cell metabolism 26, 157–170.e157. 10.1016/j.cmet.2017.06.006.

45. Netland, P.A., Scott, M.L., Boyle, J.W.t., and Lauderdale, J.D. (2011). Ocular and systemic findings in a survey of aniridia subjects. Journal of AAPOS: the official publication of the American Association for Pediatric Ophthalmology and Strabismus 15, 562–566. 10.1016/j.jaapos.2011.07.009.

46. Méjécase, C., Harding, P., Sarkar, H., Eintracht, J., Lima Cunha, D., Toualbi, L., and Moosajee, M. (2020). Generation of two human control iPS cell lines (UCLi016-A and UCLi017-A) from healthy donors with no known ocular conditions. Stem cell research 49, 102113. 10.1016/j.scr.2020.102113.

47. Torriano, S., Erkilic, N., Baux, D., Cereso, N., De Luca, V., Meunier, I., Moosajee, M., Roux, A.F., Hamel, C.P., and Kalatzis, V. (2018). The effect of PTC124 on choroideremia fibroblasts and iPSC-derived RPE raises considerations for therapy. Scientific reports 8, 8234. 10.1038/s41598-018-26481-7.

48. Sarkar, H., Mitsios, A., Smart, M., Skinner, J., Welch, A.A., Kalatzis, V., Coffey, P.J., Dubis, A.M., Webster, A.R., and Moosajee, M. (2019). Nonsense-mediated mRNA decay efficiency varies in choroideremia providing a target to boost small molecule therapeutics. Human molecular genetics 28, 1865–1871. 10.1093/hmg/ddz028.

49. Moosajee, M., Tracey-White, D., Smart, M., Weetall, M., Torriano, S., Kalatzis, V., da Cruz, L., Coffey, P., Webster, A.R., and Welch, E. (2016). Functional rescue of REP1 following treatment with PTC124 and novel derivative PTC-414 in human choroideremia fibroblasts and the nonsense-mediated zebrafish model. Human molecular genetics 25, 3416–3431. 10.1093/hmg/ddw184.

50. Reichman, S., and Goureau, O. (2016). Production of Retinal Cells from Confluent Human iPS Cells. Methods in molecular biology (Clifton, N.J.) 1357, 339–351. 10.1007/7651_2014_143.

