## Supplementary data for "Restoration of functional PAX6 in aniridia patient iPSC-derived ocular tissue models using repurposed nonsense suppression drugs"

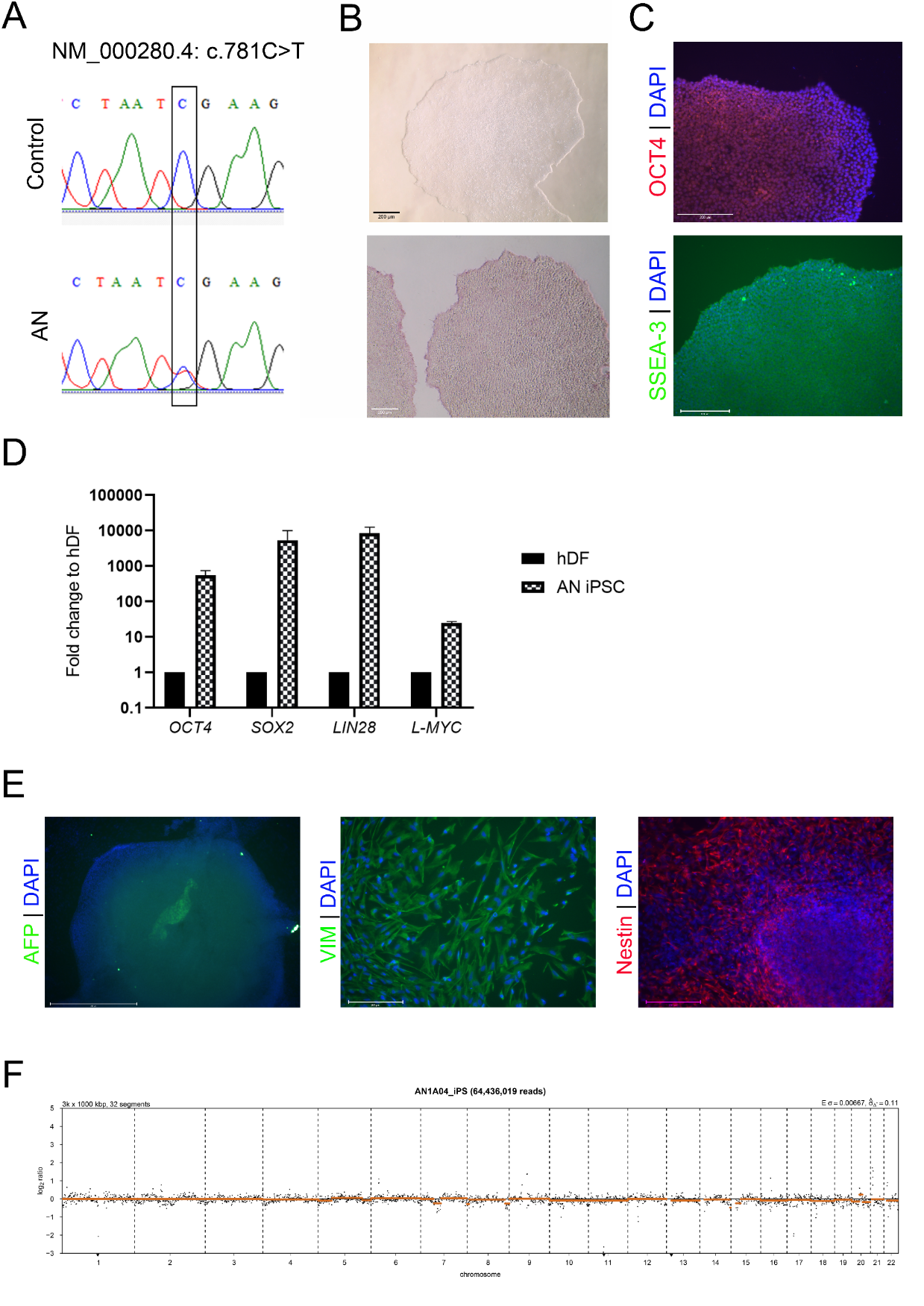
**Figure S1. Characterisation of iPSCs generated from an aniridia patient (AN) carrying a *PAX6* heterozygous nonsense variant. (A)** The heterozygous nonsense variant c.781C>T, p.Arg261* in PAX6 gene (NM_000280.4) was confirmed in aniridia iPSCs (AN) through direct sequencing and was abset from control iPSCs. **(B)** Embryonic stem cell-like morphology (upper panel) and positive alkaline phosphatase staining (lower panel) in AN iPSCs. **(C)** Positive expression of pluripotency markers OCT4 (upper panel) and SSEA-3 (lower panel). Scale bar 200µm. **(D)** Pluripotency marker genes *OCT4*, *SOX2*, *L-MYC* and *LIN28* were upregulated in AN iPSC compared to parental fibroblasts (hDF) by qRT-PCR. **(E)** In vitro differentiation ability was confirmed by random differentiation of AN iPSCs: cells stained positive for endoderm (AFP), mesoderm (Vimentin) and ectoderm (Nestin) markers. **(F)** Low-pass whole genome sequencing analysis revealed no abnormalities in AN iPSCs, showing 46,XY karyotype.

**Table S1. Primer sequences used for qRT-PCR.**

| **Marker** | **Forward sequence (5’-3’)** | **Reverse sequence (5’-3’)** | **Reference** |
| --- | --- | --- | --- |
| *GAPDH* | ACA GTT GCC ATG TAG ACC | TTT TTG GTT GAG CAC AGG | In house |
| *ACTB* | TTC TAC AAT GAG CTG CGT G | GGG GTG TTG AAG GTC TCA AA | In house |
| *PAX6* | GGC CGA ACA GAC ACA GCC CTC AC | ATC ATA ACT CCG CCC ATT CAC C | In house |
| *RAX* | AGG CGG AAA AAT AGA GTT TG | TAC CCC AAT ATT CAC TCC TC | KickStart, Sigma Aldrich |
| *VSX2* | GGC GAC ACA GGA CAA TCT TTA | TTC CGG CAG CTC CGT TTT C | KickStart, Sigma Aldrich |
| *MKi67* | AAA CCA ACA AAG AGG AAC ACA AAT T | GTC TGG AGC GCA GGG ATA TTC | In house |
| *TP63α* | ATG TCG AAA TTG CTC AGG GAT TTT CAG A | TGA CCA CCA TCT ATC AGA TTG AGC ATT ACT | Foster et al, 2019 |
| *ΔNP63* | GAA AAC AAT GCC CAG ACT CAA TTT | TCT GCG CGT GGT CTG TGT TAT | Foster et al, 2019 |
| *ABCG2* | TCC ACT GCT GTG GCA TTA AA | CCT GCT TGG AAG GCT CTA TG | Foster et al, 2019 |
| *KRT14* | CGG CCT GCT GAG ATC AAA GA | TCT GCA GAA GGA CAT TGG CA | Foster et al, 2019 |
| *SOX10* | CTC TGG AGG CTG CTG AA | TGG GCT GGT ACT TGT AGT C | Leung et al, 2016 |
